# Predicting Mycoplasma tissue and host specificity from genome sequences

**DOI:** 10.1101/2022.08.08.503189

**Authors:** Niels A. Zondervan, Vitor A. P. Martins dos Santos, Maria Suarez-Diez

## Abstract

To gain insights into the genotype-phenotype relationships in Mycoplasmas, we set to investigate which Mycoplasma proteins are most predictive of tissue and host trophism and to which functional groups of proteins they belong. We retrieved and annotated 430 *Mycoplasma* genomes and combined their genome information with data on which host and tissue these *Mycoplasmas* were isolated from. We assessed clustering of *Mycoplasma* strains from a wide range of hosts and tissues based on different functional groups of proteins. Additionally, we assessed clustering using only a subset of *M. pneumoniae* strains based on different functional groups of proteins. We found that proteins belonging to the Gene Ontology (GO) Biological process group ‘*Interspecies interaction between organisms’* proteins are most important for predicting the pathogenesis of *Mycoplasma* strains whereas for *M. pneumoniae*, those belonging to *‘Quorum sensing’* and *‘Biofilm formation’* proteins are most important for predicting pathogenesis.

Two Random Forest Classifiers were trained to accurately predicts host and tissue specificity based on only 12 proteins. For *Mycoplasma* host specificity CTP synthase complex, magnesium transporter MgtE, and glycine cleavage system are most important for correctly classifying *Mycoplasma* strains that infect humans, including opportunistic zoonotic strains. For tissue specificity, we found that a) known virulence and adhesions factor Methionine sulphate reductase MetA is predictive of urinary tract *infecting Mycoplasmas;* b) an extra cytoplasmic thiamine binding lipoprotein is most predictive of gastro-intestinal infecting *Mycoplasmas;* c) a type I restriction endonuclease is most predictive of respiratory infecting *Mycoplasmas*, and; d) a branched-chain amino acid transport system is most predictive for blood infecting *Mycoplasmas*. These findings can aid in predicting host and tissue specific pathogenicity of *Mycoplasmas* as well as provide insight in which proteins are important for specific host and tissue adaptations. Furthermore, these results underscore the usefulness of deploying genome-wide methodologies for gaining insights into pathogenicity from genome sequences.

## Introduction

*Mycoplasmas* are bacteria that adapted to live in their host environment through genome reduction, resulting in a small genome and small cell size [1], [2]. Despite their small genome size, *Mycoplasmas* still contain many genes that are not essential but enhance growth in the various host conditions encountered [3]–[5]. *Mycoplasmas* have greatly reduced metabolic capabilities growing only under the fastidious conditions of their selective host [6]. This makes it difficult to grow *Mycoplasma* on serum free defined media. For *M. pneumoniae*, a defined media has been developed by analysing their membrane components and metabolic capabilities and adding those lipids to the medium that normally are directly recruited from the host environment [7]–[9].

Because of their strong host adaptation, pathogenesis of *Mycoplasmas* is hard to typify since their ability to infect and survive in a host is largely a systems property of their obligatory pathogenic lifestyle, and not the result of a well-defined set of virulence proteins. Only few *Mycoplasma* proteins are directly categorised as pathogenic based on their GO Biological Function annotation. Examples of these proteins are *M. pnueumoniae* CARD toxins, adhesins, motility proteins and hydrogen peroxide production which are directly associated to virulence [10], [11]. These virulence proteins are however not essential, whereas many metabolic proteins such as glycerol metabolism proteins GlpF *and* GlpK are essential for *M. pneumoniae* growth in a host [12].

Previously, clustering based on functional groups of proteins was successfully used to predict pathogenic traits such as zoonotic potential for *Streptococcus suis* and *Streptococcus agalactiae* [13]. Here, we use a similar approach combined which systematic collection of meta-data from the BioSamples database [14] to identify those proteins predictive of *Mycoplasma* host and tissue infection types. The BioSample database contains tissue and host isolation data for a larger number of *Mycoplasmas* than was previously available for *Streptococcus suis* and *Streptococcus agalactiae*, allowing for better training and validation of machine learning models. We adjusted our approach to *Mycoplasmas* by selecting those functional groups of proteins known from literature to be important for pathogenesis of *Mycoplasmas*. We compared clustering of 430 *Mycoplasmas* based on 19 Gene ontology (GO) Biological process categories of proteins associated with pathogenesis to identify functional groups of proteins important for *Mycoplasmas* in general as well as *M. pneumonia* pathogenesis specifically. We combined this approach with random forest classification to accurately predicts 3 host and 4 tissue isolation sites for *Mycoplasma* genomes and to identify those proteins important for each host and tissue type.

## Materials and methods

### Genome retrieval and annotation

We retrieved 430 completely assembled Mycoplasma genomes from EBI-ENA using the Python EnaBrowserTool [15]. A list of these genomes can be found in Supplementary material 1A. Semantic Annotation Platform with Provenance (SAPP) [16] and Genome Biology Ontology Language (GBOL) [17] were used to perform *de novo* annotation and to store annotated genomes as graph files. Gene calling was performed using Prodigal 2.6.3 with codon table 4 [18]. Protein domains were identified by InterProScan 83.0 by their Pfam identifier [19]. The GNU “parallel” package version 20161222 was used to perform all the above steps in parallel [16]. Graph files were loaded in GraphDB Free version 9.7.0. Additionally, taxonomic information from UniProt was downloaded in RDF format and loaded in GraphDB. The GraphDB SPARQL endpoint was queried using the Python SPARQLWrapper [20] package and the R Curl package [21]. Genes are annotated by their protein signature, which we defined as the protein PFAM domains present in the protein. We defined such signature by ascendingly ordering all domains present in a protein concatenated with a ”;” between the different domains.

### Functional Analysis

Meta data for 430 genomes and 187 species was retrieved from the Biosamples [14] database using their API and was stored in GraphDB. The Biosample data was normalised by combining different metadata fields and standardizing the labels used for host and tissue types and was combined with taxonomic information from UniProt. Gene Ontology (GO) annotation from the GODM (GO Domain Miner) database [22] was added to protein annotation based on their domain content. The GraphDB SPARQL endpoint was queried using the Python SPARQLWrapper [20] package and the R Curl package [21] to retrieve information and store them as tab-separated files. These tab-separated files were used for all subsequent analyses. We used a literature study to identify 19 GO Biological process ontology terms with known or suspected association to pathogenesis [23]–[27].

*M. pneumoniae* genomes were overrepresented in the dataset (165 out of 430 genomes), therefore we reduced the number of *M. pneumoniae* genomes to 12 randomly selected genomes. Functional trees were build based on each of the 19 GO Biological functional groups of proteins as well as a reference tree based on all proteins. We analysed the similarity in cophenetic clustering of these 19 functional trees using the R dendextend package version 1.13.2. In addition, we repeated the clustering of these functional trees using only the 165 *Mycoplasma pneumoniae* genomes. The results of the cophenetic clustering of these functional trees based on all *Mycoplasmas* and *M. pneumoniae* were compared and plotted using the R pheatmap package version 1.0.12.

### Random Forest classification

Two Random Forest classifiers were trained using sklearn version 0.24.2 to predict host and tissue trophism of *Mycoplasma* species. The host classifier was trained to predict the three host classes: *human, pig-boar, ruminant*. The tissue isolation site classifier was trained to predict 4 isolation sites: *respiratory, blood, gastro-intestinal track*, and *uri-genital track*. The data was cleaned by removing all classes with less than 5 instances before splitting the data into 75% Training and 25% Test samples. The classifier was scored based on ‘*f1_macro’*, meaning that for each class, the performance is weighted equally irrespective of the number of samples, scoring the classifier while balancing false positives and negatives.

Hyper parameter optimization was performed using a grid search for 300 combinations of the parameters *n_estimators, max_features, max_depth, min_samples_split*, and *min_samples*. Ten times Cross validation and out of bag samples were used to avoid overfitting on the training data. The trained Random Forest classifiers were used as basis to iteratively reduce features, using Treeinterpreter 0.1.0 to interpret feature (protein) importance for overall classification as well as for specific tissue and host classes. For each iteration, the least important feature was removed until a set of the 12 most predictive features (proteins) were left. Lastly, we performed a second round of grid hyper parameter optimization when using the reduced set of protein feature. The heatmaply R package version 1.0.12 was used to plot feature importance and contribution.

## Results & Discussion

The 430 annotated genomes contained 2306 unique proteins, of which 10% were multi-domain proteins. Genomic information was merged with GO Biological function annotation for proteins and with sample isolation metadata for the 430 *Mycoplasma* genomes from the BioSamples database. Host isolation data was available for 394 genomes and tissue-isolation information was available for 157 genomes. We normalized the metadata by combining different metadata fields standardizing labels to describe the same infection host and tissue in different samples. *M. pneumoniae* was overrepresented in the original data set with 165 out the 430 genomes. Therefore, we limited the number of *M. pneumoniae* genomes to 12 randomly selected genomes in our *Mycoplasma* dataset resulting in a dataset of 277 genomes. The 165 *M. pneumoniae* genomes were kept as a separate dataset.

### Clustering based on GO functional groups of proteins

Based on literature research, we created a list of 19 GO categories expected to be associated to pathogenesis. We investigated the cophenetic distance between phylogenetic trees based on these 19 GO categories as well as a reference tree based on all proteins. The cophenetic distance is a measure of distance between genomes that have been clustered in two dendrograms providing a single similarity score between two trees. By combining all pairwise cophenetic distance scores we can create a heatmap showing the (dis)-similarity of trees based on the 19 GO functional groups of proteins (see Figure 1). A list of the 19 GO IDs with their labels can be found in Table 1.

**Figure 1.**
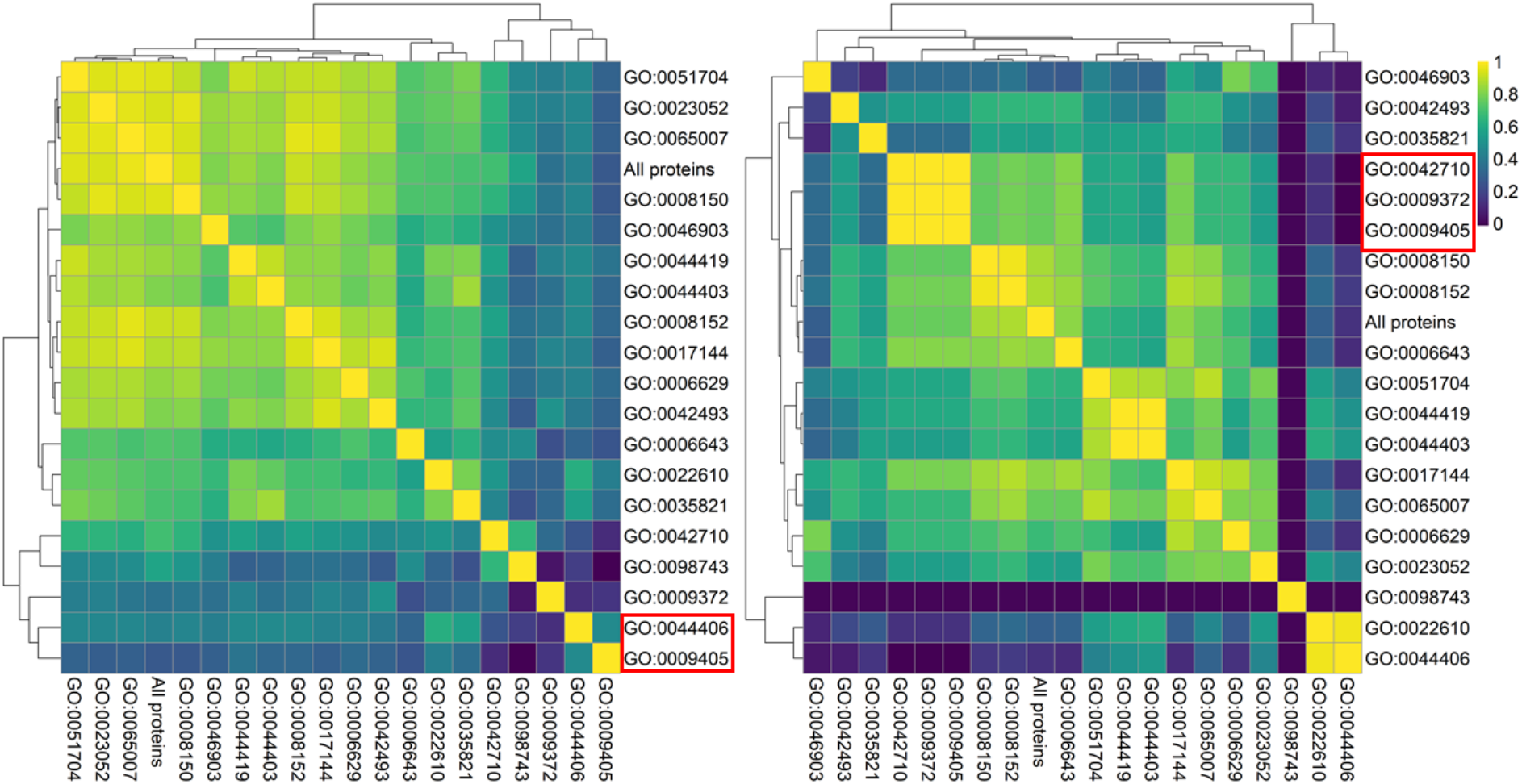
Left cophenetic correlation of GO functional trees based on All Mycoplasma. Right Cophenetic clustering based on M. pneumoniae genomes only

**Table 1.** GO ID’s associated to Pathogenesis together with their names

| GO ID | DESCRIPTION |
| --- | --- |
| GO:0008150 | Biological process |
| GO:0008152 | *Metabolic process |
| GO:0006629 | Lipid metabolic process |
| GO:0006643 | membrane lipid metabolic process |
| GO:0017144 | Drug metabolic process |
| GO:0042493 | Response to drug |
| GO:0023052 | *Signalling |
| GO:0065007 | *Biological regulation |
| GO:0022610 | *Biological adhesion |
| GO:0044406 | Adhesion of symbiont to host |
| GO:0051704 | Multi-organism process |
| GO:0044419 | Inter species interaction between organisms |
| GO:0042710 | Biofilm formation |
| GO:0098743 | Cell aggregation |
| GO:0044403 | Symbiont process |
| GO:0009372 | Quorum sensing |
| GO:0035821 | Modification of morphology or physiology of other organism |
| GO:0009405 | Pathogenesis |
| GO:0046903 | Secretion |

**Table 2.**
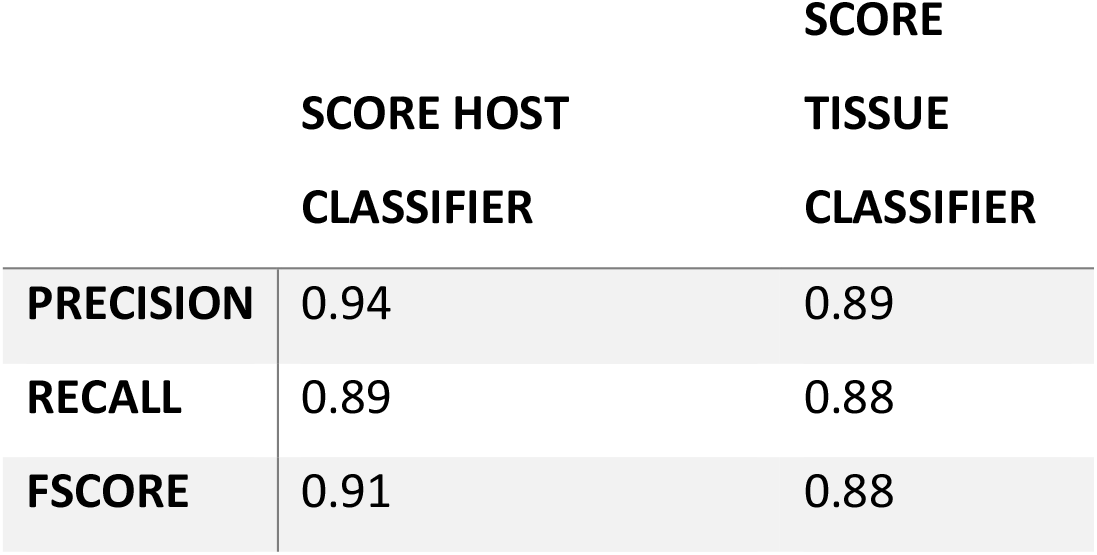
Classifier score on independent test data using the 12 most important protein features

We compared the heatmap of the similarity of these 19 trees based on the 277 *Mycoplasma* genomes with the similarity of the 19 trees based on 165 *M. pneumoniae* genomes. The objective of this comparison was to learn which GO functional groups of proteins have the highest similarity in clustering of genomes based on *‘Pathogenesis’* proteins cluster. By performing the clustering both for *Mycoplasma* genomes in general as well as for *M. pneumoniae* genomes only, we identified which functional groups of proteins are likely to be important for *Mycoplasma* pathogenesis as well as which functional groups of proteins are most likely to be important for *M. pneumoniae* pathogenesis (Figure 1).

As can be seen in Figure 1, for *Mycoplasma*, the tree based on proteins belonging to the GO category ‘*Inter species interaction between organisms’* (GO:0044419) has the highest similarity with the tree based on proteins annotated with the GO term *‘Pathogenesis’* (GO:0009405). We postulate that this high similarity might indicate that pathogenesis of *M. pneumoniae* involves proteins with functions in ‘*Symbiont or Multi interspecies interactions’*. Further investigation shows that this similarity was due to a high overlap in annotation, resulting in the same proteins being in both functional groups of proteins. All 26 *Mycoplasma* proteins annotated with the GO term *‘Pathogenesis’* contained at least one domain, which is also present in the 147 proteins annotated belonging to ‘*Interspecies interaction between organisms’*. This overlap in annotation shows that these 26 proteins from the GO Biological functional group ‘*Interspecies interaction between organisms’* is at least important for *Mycoplasma* pathogenesis.

For *M. pneumoniae*, the trees based on ‘*Quorum sensing’* (GO:0009372) and *‘Biofilm formation’* (GO:0042710) on proteins belonging to the GO category proteins have the highest similarity (1.0) as the tree based on proteins belonging to the GO category ‘*Pathogenesis*’. Not a single protein in these three categories is present in any of the other three GO categories, ruling out overlap in annotation as the reason for the high similarity of the trees. The high similarity in clustering might therefore indicate that proteins in the categories ‘*Quorum sensing’* and *‘Biofilm formation’* are important for *M. pneumoniae’s* pathogenesis.

Multiple studies support the notion that biofilm formation is important for *M. pneumoniae*’s pathogenesis. For example, Type 1 and Type 2 *M. pneumoniae* which have different phenotypes have different biofilms [28]. Biofilm formation is implicated in in chronic infections, with *M. pneumoniae* cells aggregation being important for infections [29]. We found no studies confirming the importance of ‘*Quorum sensing’* sensing for *M. pneumoniae* pathogenesis. However, it would not come as a surprise if Quorum sensing would be important for *M. pneumoniae* pathogenesis since quorum sensing plays an important role in pathogenesis of other lung infecting bacteria such as *Streptococcus pneumoniae* [30] and *Klebsiella pneumoniae* by regulating virulence systems such as ESX-3, biofilm formation, and secretion of PgaA porin [31], [32]. *M. pneumoniae* virulence systems such as CARD toxins are upregulated when in contact with host cells and in acidic conditions. We postulate that quorum sensing proteins are likely to be involved in pathogenesis by for example sensing host conditions, cell to cell contact to regulate motility, and virulence.

### Zoonotic potential

Only two zoonotic strains were identified in our dataset, GCA:001005165 and GCA:000012765, belonging to the *M. capricolum* species group. Two out of the 13 strains were isolated from humans, 2 from goats and 1 from a Tibetan Antilope. The remaining 8 *M. capricolum* genomes have no host isolation data available. We see that *M. capricolum* is taxonomically mixed with *M. leachii* which infects cow. The other M. capricolum isolates are mostly from the respiratory tract, while one zoonotic strain was found in the bloodstream. For the other, no tissue isolation data is available. We hypothesize that *M. capricolum* zoonotic capability is likely limited to infecting the blood stream. In general, it appears that *Mycoplasma* species are so adjusted to their host that they have a limited zoonotic potential [1].

### Predicting host and tissue trophism

Two random forest classifiers were built to predict *Mycoplasma* strains host and tissue infection site respectively. The data was filtered on host classes with a minimum of 5 strains associated to them, resulting in 125 genomes and 3 classes for the host classifier: *human, pig-boar, ruminant*.

Similarly, data was filtered on tissue classes with a minimum of 5 strains associated to them, resulting in 91 genomes and 4 classes for the tissue classifier: *blood, gastro-intestinal, respiratory, and uri-genital tracks*.

The resulting datasets were separated in 75% training and 25% test data. Models were fitted with all protein features using hyper parameter tuning followed by iterative feature reduction to select a set of the 12 most important protein features for classification. A final round of hyper parameter optimization was performed. Models were trained giving equal weight to each class to consider unequal numbers of the classes in both training and test data.

The resulting classifiers predict host and tissue specificity with a high precision on independent dataset not used to train the classifiers (Table1A). An overview of optimal hyper parameters as well as the scores for the classifier for both the full and the reduced set of features 12 features, and confusion matrixes can be found in Supplementary file 1B.

The classifiers predict the provided classes with great accuracy, precision, and recall using only 12 features. The 12 protein features were analysed for their contribution and overall importance for classification. No overlap between the features used by these two classifiers was observed. For the host classifier, we see that more protein features are predictive of a single isolation host type. For the tissue classifier, we see that more protein features are more synergistic in their prediction, being associated to multiple tissue isolation sites.

#### Host classification

The host classifier using >2000 features performed only slightly better with a *f1_score* of 0.97 versus 0.94 using the most 12 important features (Supplementary material 1B). A slightly lower *f1_score* for the test than training data is to be expected since it is likely that some rarely occurring proteins only occur in the training data and not in the test data. Inspection of the confusion matrix (Supplementary material 1B) revealed one strain isolated from a human and one strain isolated from a *pig-boar* genome to be wrongly classified. One of these genomes, GCA_001005165, belongs to a zoonotic *Mycoplasma capricolum* isolated from human blood which was misclassified as being isolated from a ruminant. The misclassification as well as the absence of any clear difference in its pathogenesis proteins from other *M. capricolum* strains indicates this zoonotic potential might be the result of opportunism. The other of the two misclassified genome is GCA_000815065 from *M. flocculare*, which was wrongly classified as being isolated from a ruminant.

#### Tissue classification

The tissue isolation site classifier using all >2000 features performed only slightly better with a *f1_score* of the 0.89 versus an *f1_score* of 0.84 when only using the 12 most important features. Only three genomes (GCA_000319465, GCA_012934855 and GCA_017389835) were misclassified based on the classifier with the 12 most important features. The first genome is from *Mycoplasma haemominutum ‘Birmingham 1’;* the second genome is from *M. phocoena* infects the urinary tract of harbour porpoise; and the third genome is from an unspecified *Mycoplasma* from the gut microbiome of a buffalo. The first two genomes are from species that only occur 1 time in our dataset while the third genome turned out to be from the gut microbiome and not to be associated to an infection of the gut. Although the three examples above show that tissue isolation site for some genomes of rarely occurring host are misclassified, we do see that the tissue isolation site for other rarely occurring hosts in our dataset, such as one dog (GCA_000238995) and two cats (GCA_000186985, GCA_000200735) infecting *Mycoplasma*, where accurately predicted to be blood infecting. Therefore, we can conclude that tissue classification is to some extent host specific and to some extent non-host specific.

As can be seen in Figure 2, for isolation-host classification, *human* and *pig-boar* appear to have a higher similarity in their classification while for isolation-tissue, *blood* and *uri-genital* are closest.The *gastro-intestinal* classification is somewhat more dissimilar, while classification of the *respiratory tract* as isolation site is most dissimilar to the other classes.

**Figure 2.**
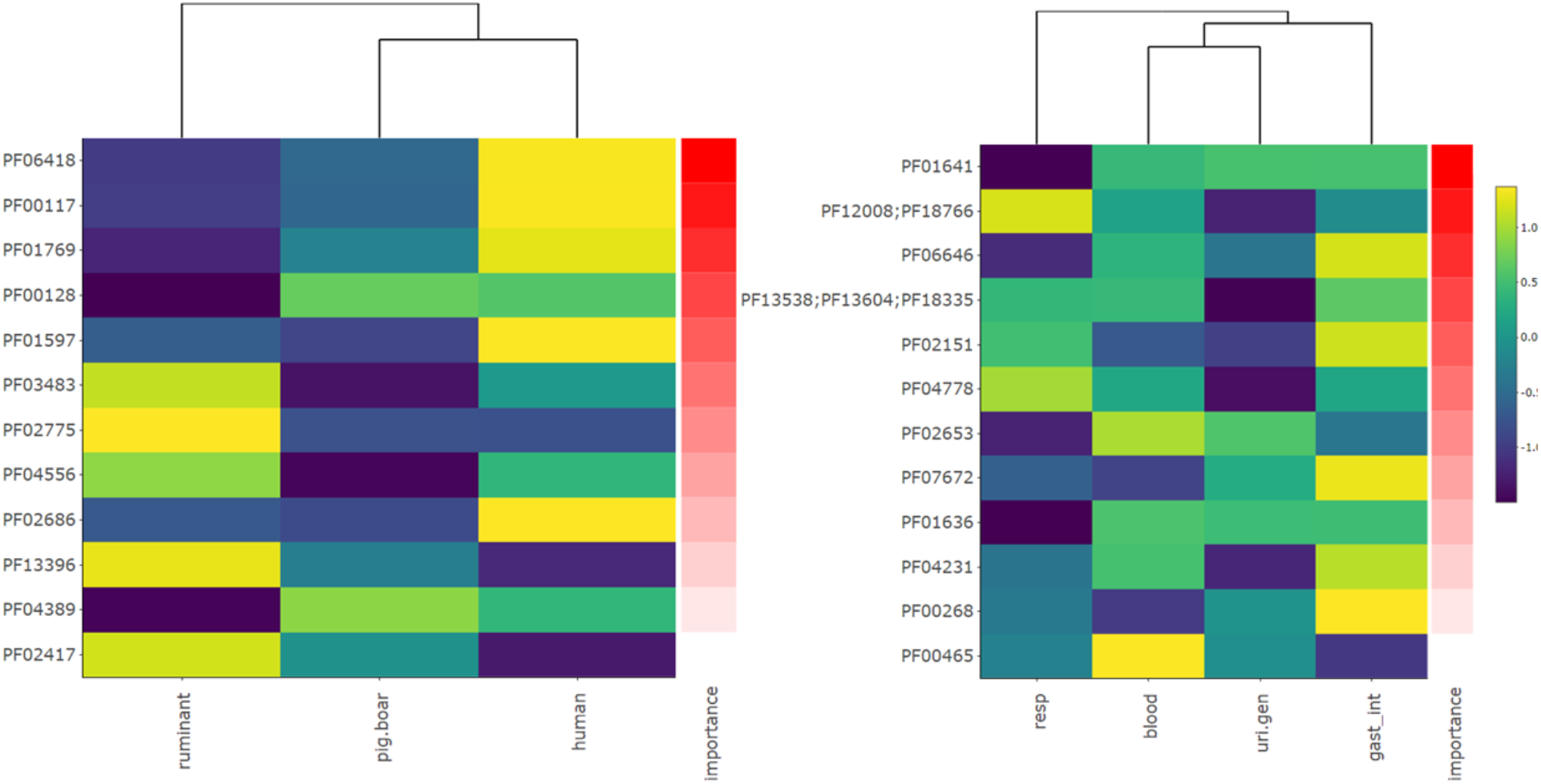
Left host isolation-site classifier feature importance and feature contribution scaled per row. Right tissue isolation-site classifier feature importance and feature contribution scaled per row.

#### Proteins important for predicting host trophism

We further investigated which protein signatures are most important for host and tissue classification for our classifiers based on the 12 most important protein features.

The Pfam [19] domains PF06418 CTP_synth_N as well as PF00117 GATase strongly contribute to predicting *human* as isolation host class. Surprisingly although very important for correctly predicting human infecting *Mycoplasma* species, the protein is only present in a few human infecting strains such as *M. capricolum* (GCA_001005165), *Candidatus Mycoplasma girerdii* (GCA_002215425), *M. penetrans* (GCA_000011225, GCA_004127945) and *Candidatus M. haemohominis* (GCA_008326325). From the 128 genomes that contain these proteins, only the 5 were isolated from humans. Among these are the 2 *M. capricolum* zoonotic strains and of the four known isolation sites, 4 were isolated from human blood infections. As such, we postulate this protein complex as a requirement for opportunistic infection of humans through the blood stream. Nucleotide synthesis is known to be critical for growth of bacteria in human blood [34].

PF01769, the magnesium transporter MgtE, is the third most important feature for host classification. This magnesium transporter has a high contribution to predicting the host class *human* and to lesser extentcontributes to predicting *pig-boar* as host class. Also, this protein is only present in three human infecting Mycoplasma, namely *Candidatus M. haemohominis* (GCA_008326325) and *M. capricolum subsp. Capricolum* (GCA_001005165) while being abundant in *pig-boar* and *ruminant* infecting *Mycoplasmas*. Although KO studies associate this protein to be magnesium transport, it is unknown if this is the primary function of this protein [11]. MgtE is involved in regulating many virulence factors in *Aeromonas hydrophila* as well as in fine tuning regulation of virulence proteins in *Pseudomonas aeruginosa* [35], [36]. PF01597, a glycine cleavage system is another important contributor to predicting human as the isolation host. This protein is however only present in a single human infecting *Mycoplasmas*, namely *M*.*capricolum subsp. Capricolum* (GCA_001005165) and was reported by Kaminga et al. [5] as a predictor of ruminant and pig infecting *Mycoplasma*. Indeed, PF01597 is much more common in *M. hyopneumoniae* but might contribute to *M. capricolum subsp. Capricolum’s* ability to opportunistically infect humans through the blood stream. As can be seen from the above examples, protein features with a high contribution to predicting a single class are not always the most commonly occurring within that class. In the case of *human* infecting *Mycoplasmas*, it appears that the most important protein features are those few proteins that help identify the few zoonotic and opportunistic *Mycoplasma* strains that infect the blood stream.

PF00128 and PF04389 have the highest contribution to predicting *pig-boar* as isolation host. P00128 is an Alpha-amylase while PF04389 is a Peptidase family M28 protein. PF00128 and PF04389 are both present in present in all *M. hyosynoviae* strains proteins. PF03483 and PF02775 have the strongest contribution to predicting *Mycoplasma* that infect ruminants. PF03483 is a B3/B4 domain found in tRNA synthetase beta subunits and other synthetases, while PF02775 is a thiamine pyrophosphate (vitamin B1) binding domain. Also, DpnII restriction endonuclease PF04556 and phospholipase_D-nuclease N-terminal PF13396 strongly contribute to predicting ruminant infecting *Mycoplasma*.

#### Proteins important for predicting tissue trophism

We further investigated proteins that are most important for tissue classification. The tissue classifiers shows that methionine sulfate reductase A (MetA) PF01641 to be important for urinary tract and genital infections. MetA is a known virulence determinant for *M. genitalium* which infects the urinary tract while being necessary for proper adhesion [37]. The second most important protein for classification is PF12008; PF18766, a type I restriction endonuclease, which has the highest contribution to predicting the class ‘*respiratory tract*’ as isolation site.

PF06646 MG289, an extra cytoplasmic thiamine binding lipoprotein has the highest contribution to predicting the *gastro-intestinal* tract as tissue isolation site, as well as its high contribution predicting *blood* as the tissue isolation site. Additionally, it was shown that MG289 enhances microbial invasion and persistence in *Mycoplasma genitalium* [38].

PF02653, a branched-chain amino acid transport system was found to have the highest contribution to classifying the tissue isolation site *blood* and the tissue isolation sites *uri-genital*. Literature confirms that transport and uptake of branched-chain amino acids is important for protein synthesis and their requirement for environmental adaptation[39]. Other obligatory parasites like the intracellular pathogen *Francisella* lost all branched-chain amino acid biosynthetic pathways and rely on dedicated uptake systems for their survival in the host [40]. Branched-chain amino acids are essential for lymphocyte responsiveness and proper functioning of other immune cells [41], which puts them at the interface of pathogen-host interaction.

#### Strengths and weaknesses of classification

We repeated the host classification, allowing genomes from rarely occurring hosts isolated from other host to be in the train and testing dataset. Although the accuracy of the training data remained high at 94%, the accuracy for the test data dropped to around 75% and the *f1_score* dropped to 55%. This means that some *Mycoplasma* from host classes contain some of the 12 important features, resulting in misclassification. Classification for strains with rarely occurring hosts is not feasible since there are too few samples for both test and training data.

We also tested if combined prediction of host and tissue was possible to find *host_tissue* specific features. Unfortunately, too few genomes remain when combining tissue and host isolation site information and filtering on a minimum of 5 genomes (Supplementary material 1B). The BioSamples metadata used in our analysis required a manual normalization, combining different metadata fields and normalizing the various labels used in these metadata fields. Therefore, we want to emphasize the importance of 1) sequencing larger numbers of genomes, also for more rare hosts and 2) the importance of well-defined metadata for sequenced genomes to allow for machine learning approaches to predict strain specific properties such as the host isolation and tissue isolation site.

The use of these host and tissue classifiers reveals some of the strengths and weaknesses of classification. For example, those proteins identified as most important for classifications of *human* are those which can best predict the few strains that infect *human* instead of their normal host. Furthermore, because we minimize the number of proteins needed for our prediction, we find proteins that best split species over multiple classes, meaning that many proteins that are identified as important for classification are present in multiple host or tissue types. This makes the approach used in this study less suitable for identifying proteins typical for a single host or tissue type. Furthermore, it should be noted that our classifiers are optimized to predict all available classes equally optimizing for the *f1_score* to balance false positives and false negatives. This is desirable when creating a robust classifier without favouring any single class as intended in our study. However, for medical purposes, such as predicting the risk of a zoonotic outbreak, having maximum recall for *human* infecting *Mycoplasmas* would be desirable since false positives would be preferred over false negatives. Therefore, we advise to train new classifiers when used for such purposes.

## Conclusion

In this study we demonstrated that clustering based on different GO Biological function categories of proteins for *M. pneumoniae*, can provide insight in terms of which functional groups of proteins are most important for pathogenesis. Our study revealed differences in the functional groups of proteins important for *M. pneumoniae* pathogenesis and the pathogenesis of *Mycoplasmas* in general. The GO functional group of proteins *Interspecies interaction between organisms’* is important for pathogenesis of *Mycoplasmas* in general, while ‘*Quorum sensing’* and *‘Biofilm formation’* proteins are important for *M. pneumoniae* pathogenesis.

Furthermore, we show that a small set of proteins can be used to classify host and tissue specificity of various *Mycoplasmas*. Most proteins important for classification were found to have corroborative evidence for their importance to be available in literature, while some might provide new insights such as the proteins identified that differentiate human infecting zoonotic strains from non-zoonotic strains. Finally, our analyses show the feasibility of predicting species properties such as host and tissue types based on genomic information, as well as the importance of high-quality sample meta-data to enable classification through machine learning.

## Supporting information

Genomes and supplementary figures

## Declarations

## Abbreviations

GO: Genome Ontology
t-SNE: T-distributed Stochastic Neighbour Embedding

## Availability of data and materials

The authors declare that all data supporting the findings of this study are available within the article and its supplementary information.

## Author contribution

NZ performed the main analyses and wrote the draft manuscript. MSD and VDMS participated in the design of the study and supervised and directed the research. MSD and VDMS revised the manuscript. All authors contributed to the writing of the final version of the manuscript.

## Supplementary material

Additional file 1AB List of Mycoplasma genomes & Random Forest classification of host and tissue isolation data

