## Supplementary material for "Predicting Mycoplasma tissue and host specificity from genome sequences": Genomes and supplementary figures

### Supplementary material 1A Genomes

Below a list of all genomes used in this analysis

| genbank | name | taxonID |
|---|---|---|
| GCA_000026765 | Mycoplasma conjunctivae | 45361 |
| GCA_000433455 | Mycoplasma sp. CAG:877 | 1262907 |
| GCA_000433475 | Mycoplasma sp. CAG:956 | 1262908 |
| GCA_000434175 | Mycoplasma sp. CAG:611 | 1262905 |
| GCA_000437315 | Mycoplasma sp. CAG:472 | 1262904 |
| GCA_000438195 | Mycoplasma sp. CAG:776 | 1262906 |
| GCA_000691265 | Mycoplasma hyosynoviae | 29559 |
| GCA_000691285 | Mycoplasma hyosynoviae | 29559 |
| GCA_000691305 | Mycoplasma hyosynoviae | 29559 |
| GCA_000691325 | Mycoplasma hyosynoviae | 29559 |
| GCA_000691345 | Mycoplasma hyosynoviae | 29559 |
| GCA_000691365 | Mycoplasma hyosynoviae | 29559 |
| GCA_000691385 | Mycoplasma hyosynoviae | 29559 |
| GCA_000733865 | Mycoplasma buteonis | 171280 |
| GCA_000759375 | Mycoplasma hominis | 2098 |
| GCA_000759385 | Mycoplasma hominis | 2098 |
| GCA_000770195 | Candidatus Mycoplasma girerdii | 1318617 |
| GCA_000828855 | Mycoplasma canadense | 29554 |
| GCA_000935865 | Mycoplasma hominis | 2098 |
| GCA_000935875 | Mycoplasma hominis | 2098 |
| GCA_000941075 | Mycoplasma dispar | 86660 |
| GCA_000947915 | Mycoplasma hominis | 2098 |
| GCA_001017595 | Mycoplasma hominis | 2098 |
| GCA_001063275 | Mycoplasma hominis | 2098 |
| GCA_001063305 | Mycoplasma hominis | 2098 |
| GCA_001199495 | Mycoplasma sp. HU2014 | 1664275 |
| GCA_001272875 | Mycoplasma pneumoniae | 2104 |
| GCA_001296485 | Mycoplasma pneumoniae | 2104 |
| GCA_001296505 | Mycoplasma pneumoniae | 2104 |
| GCA_001296515 | Mycoplasma pneumoniae | 2104 |
| GCA_001296525 | Mycoplasma pneumoniae | 2104 |
| GCA_001296565 | Mycoplasma pneumoniae | 2104 |
| GCA_001296585 | Mycoplasma pneumoniae | 2104 |
| GCA_001296605 | Mycoplasma pneumoniae | 2104 |
| GCA_001296615 | Mycoplasma pneumoniae | 2104 |
| GCA_001296625 | Mycoplasma pneumoniae | 2104 |
| GCA_001296665 | Mycoplasma pneumoniae | 2104 |
| GCA_001296685 | Mycoplasma pneumoniae | 2104 |

| | | |
|---|---|---|
| *GCA_001296705* | Mycoplasma pneumoniae | 2104 |
| *GCA_001296725* | Mycoplasma pneumoniae | 2104 |
| *GCA_001296735* | Mycoplasma pneumoniae | 2104 |
| *GCA_001296765* | Mycoplasma pneumoniae | 2104 |
| *GCA_001296785* | Mycoplasma pneumoniae | 2104 |
| *GCA_001296805* | Mycoplasma pneumoniae | 2104 |
| *GCA_001296815* | Mycoplasma pneumoniae | 2104 |
| *GCA_001296825* | Mycoplasma pneumoniae | 2104 |
| *GCA_001296855* | Mycoplasma pneumoniae | 2104 |
| *GCA_001296885* | Mycoplasma pneumoniae | 2104 |
| *GCA_001296895* | Mycoplasma pneumoniae | 2104 |
| *GCA_001296905* | Mycoplasma pneumoniae | 2104 |
| *GCA_001455605* | Mycoplasma pneumoniae | 2104 |
| *GCA_001455625* | Mycoplasma pneumoniae | 2104 |
| *GCA_001455635* | Mycoplasma pneumoniae | 2104 |
| *GCA_001455675* | Mycoplasma pneumoniae | 2104 |
| *GCA_001455685* | Mycoplasma pneumoniae | 2104 |
| *GCA_001455695* | Mycoplasma pneumoniae | 2104 |
| *GCA_001455735* | Mycoplasma pneumoniae | 2104 |
| *GCA_001455775* | Mycoplasma pneumoniae | 2104 |
| *GCA_001455795* | Mycoplasma pneumoniae | 2104 |
| *GCA_001509195* | Mycoplasma pneumoniae | 2104 |
| *GCA_001558175* | Mycoplasma pneumoniae | 2104 |
| *GCA_001641205* | Mycoplasma sp. Bg1 | 1784822 |
| *GCA_001641225* | Mycoplasma sp. Bg2 | 1784823 |
| *GCA_001645765* | Candidatus Mycoplasma haemobos | 432608 |
| *GCA_001676495* | Mycoplasma gallisepticum | 2096 |
| *GCA_001676505* | Mycoplasma gallisepticum | 2096 |
| *GCA_001676515* | Mycoplasma gallisepticum | 2096 |
| *GCA_001676525* | Mycoplasma gallisepticum | 2096 |
| *GCA_001676575* | Mycoplasma gallisepticum | 2096 |
| *GCA_001683635* | Mycoplasma gallisepticum | 2096 |
| *GCA_001683675* | Mycoplasma gallisepticum | 2096 |
| *GCA_001705605* | Mycoplasma hyorhinis | 2100 |
| *GCA_001705745* | Mycoplasma gallisepticum | 2096 |
| *GCA_001714895* | Mycoplasma ovipneumoniae | 29562 |
| *GCA_001715025* | Mycoplasma ovipneumoniae | 29562 |
| *GCA_001715035* | Mycoplasma ovipneumoniae | 29562 |
| *GCA_001715045* | Mycoplasma ovipneumoniae | 29562 |
| *GCA_001715095* | Mycoplasma ovipneumoniae | 29562 |
| *GCA_001715105* | Mycoplasma ovipneumoniae | 29562 |
| *GCA_001715115* | Mycoplasma ovipneumoniae | 29562 |
| *GCA_001901705* | Mycoplasma pneumoniae | 2104 |
| *GCA_001906845* | Mycoplasma hominis | 2098 |

| | | |
|---|---|---|
| *GCA_001906855* | Mycoplasma hominis | 2098 |
| *GCA_001917275* | Mycoplasma sp. CAG:611_25_7 | 1897009 |
| *GCA_002090215* | Mycoplasma pneumoniae | 2104 |
| *GCA_002090235* | Mycoplasma pneumoniae | 2104 |
| *GCA_002090275* | Mycoplasma pneumoniae | 2104 |
| *GCA_002090295* | Mycoplasma pneumoniae | 2104 |
| *GCA_002090315* | Mycoplasma pneumoniae | 2104 |
| *GCA_002095995* | Mycoplasma pneumoniae | 2104 |
| *GCA_002096015* | Mycoplasma pneumoniae | 2104 |
| *GCA_002096035* | Mycoplasma pneumoniae | 2104 |
| *GCA_002127985* | Mycoplasma pneumoniae | 2104 |
| *GCA_002128005* | Mycoplasma pneumoniae | 2104 |
| *GCA_002128025* | Mycoplasma pneumoniae | 2104 |
| *GCA_002128045* | Mycoplasma pneumoniae | 2104 |
| *GCA_002128065* | Mycoplasma pneumoniae | 2104 |
| *GCA_002128085* | Mycoplasma pneumoniae | 2104 |
| *GCA_002128105* | Mycoplasma pneumoniae | 2104 |
| *GCA_002128125* | Mycoplasma pneumoniae | 2104 |
| *GCA_002128145* | Mycoplasma pneumoniae | 2104 |
| *GCA_002128165* | Mycoplasma pneumoniae | 2104 |
| *GCA_002128185* | Mycoplasma pneumoniae | 2104 |
| *GCA_002128205* | Mycoplasma pneumoniae | 2104 |
| *GCA_002128235* | Mycoplasma pneumoniae | 2104 |
| *GCA_002128265* | Mycoplasma pneumoniae | 2104 |
| *GCA_002128285* | Mycoplasma pneumoniae | 2104 |
| *GCA_002147855* | Mycoplasma pneumoniae | 2104 |
| *GCA_002193015* | Mycoplasma hyopneumoniae | 2099 |
| *GCA_002213485* | Mycoplasma hyopneumoniae | 2099 |
| *GCA_002214445* | Mycoplasma hyosynoviae | 29559 |
| *GCA_002215425* | Candidatus Mycoplasma girerdii | 1318617 |
| *GCA_002245785* | Mycoplasma testudineum | 244584 |
| *GCA_002257505* | Mycoplasma hyopneumoniae | 2099 |
| *GCA_002298905* | Mycoplasma sp. UBA710 | 1946970 |
| *GCA_002355695* | Mycoplasma pneumoniae | 2104 |
| *GCA_002355715* | Mycoplasma pneumoniae | 2104 |
| *GCA_002563345* | Mycoplasma pneumoniae | 2104 |
| *GCA_002563355* | Mycoplasma pneumoniae | 2104 |
| *GCA_002563365* | Mycoplasma pneumoniae | 2104 |
| *GCA_002563415* | Mycoplasma pneumoniae | 2104 |
| *GCA_002563435* | Mycoplasma pneumoniae | 2104 |
| *GCA_002563495* | Mycoplasma pneumoniae | 2104 |
| *GCA_002563515* | Mycoplasma pneumoniae | 2104 |
| *GCA_002563545* | Mycoplasma pneumoniae | 2104 |
| *GCA_002736285* | Mycoplasma dispar | 86660 |

| | | |
|---|---|---|
| *GCA_002952835* | Mycoplasma hominis | 2098 |
| *GCA_003208575* | Mycoplasma alkalescens | 45363 |
| *GCA_003253435* | Mycoplasma auris | 51363 |
| *GCA_003265155* | Mycoplasma wenyonii | 65123 |
| *GCA_003269445* | Mycoplasma cloacale | 92401 |
| *GCA_003285045* | Mycoplasma anseris | 92400 |
| *GCA_003287375* | Mycoplasma hominis | 2098 |
| *GCA_003298555* | Mycoplasma hominis | 2098 |
| *GCA_003298565* | Mycoplasma hominis | 2098 |
| *GCA_003298635* | Mycoplasma hominis | 2098 |
| *GCA_003326045* | Mycoplasma hominis | 2098 |
| *GCA_003326075* | Mycoplasma hominis | 2098 |
| *GCA_003326095* | Mycoplasma hominis | 2098 |
| *GCA_003326125* | Mycoplasma hominis | 2098 |
| *GCA_003332325* | Mycoplasma phocidae | 142651 |
| *GCA_003383595* | Mycoplasma phocicerebrale | 142649 |
| *GCA_003663725* | Mycoplasma hominis | 2098 |
| *GCA_003688445* | Mycoplasma subdolum | 92407 |
| *GCA_003855455* | Mycoplasma struthionis | 538220 |
| *GCA_004011945* | Mycoplasma sp. ATU-Cv-508 | 2048001 |
| *GCA_004011965* | Mycoplasma sp. ATU-Cv-703 | 2498595 |
| *GCA_004127945* | Mycoplasma penetrans | 28227 |
| *GCA_004335975* | Mycoplasma marinum | 1937190 |
| *GCA_004335995* | Mycoplasma todarodis | 1937191 |
| *GCA_004362335* | Mycoplasma testudineum | 244584 |
| *GCA_004365165* | Mycoplasma hyosynoviae | 29559 |
| *GCA_004768725* | Mycoplasma hyopneumoniae | 2099 |
| *GCA_004771095* | Mycoplasma gallisepticum | 2096 |
| *GCA_004771115* | Mycoplasma gallisepticum | 2096 |
| *GCA_006228185* | Mycoplasma nasistruthionis | 353852 |
| *GCA_006385185* | Mycoplasma equirhinis | 92402 |
| *GCA_006385795* | Mycoplasma falconis | 92403 |
| *GCA_006491995* | Mycoplasma neophronis | 872983 |
| *GCA_006494695* | Mycoplasma nasistruthionis | 353852 |
| *GCA_007858495* | Mycoplasma anserisalpingitidis | 519450 |
| *GCA_007858515* | Mycoplasma anserisalpingitidis | 519450 |
| *GCA_007859615* | Mycoplasma anserisalpingitidis | 519450 |
| *GCA_007923985* | Mycoplasma hyopneumoniae | 2099 |
| *GCA_007924005* | Mycoplasma hyorhinis | 2100 |
| *GCA_008326325* | Candidatus Mycoplasma haemohominis | 1494318 |
| *GCA_008728895* | Mycoplasma gallisepticum | 2096 |
| *GCA_008728915* | Mycoplasma gallisepticum | 2096 |
| *GCA_008728935* | Mycoplasma gallisepticum | 2096 |
| *GCA_009664285* | Mycoplasma hominis | 2098 |

| | | |
|---|---|---|
| *GCA_009664325* | Mycoplasma hominis | 2098 |
| *GCA_009756935* | Mycoplasma ovipneumoniae | 29562 |
| *GCA_009792315* | Mycoplasma sp. NEAQ87857 | 2683967 |
| *GCA_009809995* | Mycoplasma pneumoniae | 2104 |
| *GCA_009810015* | Mycoplasma pneumoniae | 2104 |
| *GCA_009810035* | Mycoplasma pneumoniae | 2104 |
| *GCA_009810055* | Mycoplasma pneumoniae | 2104 |
| *GCA_009810075* | Mycoplasma pneumoniae | 2104 |
| *GCA_009810095* | Mycoplasma pneumoniae | 2104 |
| *GCA_009810115* | Mycoplasma pneumoniae | 2104 |
| *GCA_009810135* | Mycoplasma pneumoniae | 2104 |
| *GCA_009810155* | Mycoplasma pneumoniae | 2104 |
| *GCA_009810175* | Mycoplasma pneumoniae | 2104 |
| *GCA_009810195* | Mycoplasma pneumoniae | 2104 |
| *GCA_009810235* | Mycoplasma pneumoniae | 2104 |
| *GCA_009810255* | Mycoplasma pneumoniae | 2104 |
| *GCA_009810275* | Mycoplasma pneumoniae | 2104 |
| *GCA_009810295* | Mycoplasma pneumoniae | 2104 |
| *GCA_009810315* | Mycoplasma pneumoniae | 2104 |
| *GCA_009810335* | Mycoplasma pneumoniae | 2104 |
| *GCA_009810355* | Mycoplasma pneumoniae | 2104 |
| *GCA_009810375* | Mycoplasma pneumoniae | 2104 |
| *GCA_009810395* | Mycoplasma pneumoniae | 2104 |
| *GCA_009810415* | Mycoplasma pneumoniae | 2104 |
| *GCA_009810435* | Mycoplasma pneumoniae | 2104 |
| *GCA_009810455* | Mycoplasma pneumoniae | 2104 |
| *GCA_009810475* | Mycoplasma pneumoniae | 2104 |
| *GCA_009810495* | Mycoplasma pneumoniae | 2104 |
| *GCA_009810515* | Mycoplasma pneumoniae | 2104 |
| *GCA_009810535* | Mycoplasma pneumoniae | 2104 |
| *GCA_009810555* | Mycoplasma pneumoniae | 2104 |
| *GCA_009810595* | Mycoplasma pneumoniae | 2104 |
| *GCA_009810615* | Mycoplasma pneumoniae | 2104 |
| *GCA_009810635* | Mycoplasma pneumoniae | 2104 |
| *GCA_009810655* | Mycoplasma pneumoniae | 2104 |
| *GCA_009810675* | Mycoplasma pneumoniae | 2104 |
| *GCA_009810695* | Mycoplasma pneumoniae | 2104 |
| *GCA_009810715* | Mycoplasma pneumoniae | 2104 |
| *GCA_009810735* | Mycoplasma pneumoniae | 2104 |
| *GCA_009810755* | Mycoplasma pneumoniae | 2104 |
| *GCA_009810775* | Mycoplasma pneumoniae | 2104 |
| *GCA_009810795* | Mycoplasma pneumoniae | 2104 |
| *GCA_009810815* | Mycoplasma pneumoniae | 2104 |
| *GCA_009810835* | Mycoplasma pneumoniae | 2104 |

| | | |
|---|---|---|
| *GCA_009810855* | Mycoplasma pneumoniae | 2104 |
| *GCA_009810875* | Mycoplasma pneumoniae | 2104 |
| *GCA_009810895* | Mycoplasma pneumoniae | 2104 |
| *GCA_009810915* | Mycoplasma pneumoniae | 2104 |
| *GCA_009810935* | Mycoplasma pneumoniae | 2104 |
| *GCA_009810955* | Mycoplasma pneumoniae | 2104 |
| *GCA_009810975* | Mycoplasma pneumoniae | 2104 |
| *GCA_009810995* | Mycoplasma pneumoniae | 2104 |
| *GCA_009811015* | Mycoplasma pneumoniae | 2104 |
| *GCA_009811035* | Mycoplasma pneumoniae | 2104 |
| *GCA_009811055* | Mycoplasma pneumoniae | 2104 |
| *GCA_009811075* | Mycoplasma pneumoniae | 2104 |
| *GCA_009811095* | Mycoplasma pneumoniae | 2104 |
| *GCA_009811115* | Mycoplasma pneumoniae | 2104 |
| *GCA_009811135* | Mycoplasma pneumoniae | 2104 |
| *GCA_009811155* | Mycoplasma pneumoniae | 2104 |
| *GCA_009811175* | Mycoplasma pneumoniae | 2104 |
| *GCA_009811195* | Mycoplasma pneumoniae | 2104 |
| *GCA_009811215* | Mycoplasma pneumoniae | 2104 |
| *GCA_009811235* | Mycoplasma pneumoniae | 2104 |
| *GCA_009811255* | Mycoplasma pneumoniae | 2104 |
| *GCA_009811275* | Mycoplasma pneumoniae | 2104 |
| *GCA_009811295* | Mycoplasma pneumoniae | 2104 |
| *GCA_009939745* | Mycoplasma pneumoniae | 2104 |
| *GCA_009939765* | Mycoplasma pneumoniae | 2104 |
| *GCA_009939785* | Mycoplasma pneumoniae | 2104 |
| *GCA_009939805* | Mycoplasma pneumoniae | 2104 |
| *GCA_009939825* | Mycoplasma pneumoniae | 2104 |
| *GCA_009939845* | Mycoplasma pneumoniae | 2104 |
| *GCA_009939975* | Mycoplasma pneumoniae | 2104 |
| *GCA_009940325* | Mycoplasma pneumoniae | 2104 |
| *GCA_009940965* | Mycoplasma pneumoniae | 2104 |
| *GCA_009941325* | Mycoplasma pneumoniae | 2104 |
| *GCA_009941705* | Mycoplasma pneumoniae | 2104 |
| *GCA_009942155* | Mycoplasma pneumoniae | 2104 |
| *GCA_009942395* | Mycoplasma pneumoniae | 2104 |
| *GCA_009942655* | Mycoplasma pneumoniae | 2104 |
| *GCA_009942915* | Mycoplasma pneumoniae | 2104 |
| *GCA_009943205* | Mycoplasma pneumoniae | 2104 |
| *GCA_009943505* | Mycoplasma pneumoniae | 2104 |
| *GCA_009943805* | Mycoplasma pneumoniae | 2104 |
| *GCA_009944075* | Mycoplasma pneumoniae | 2104 |
| *GCA_009944335* | Mycoplasma pneumoniae | 2104 |
| *GCA_009944725* | Mycoplasma pneumoniae | 2104 |

| | | |
|---|---|---|
| *GCA_009945165* | Mycoplasma pneumoniae | 2104 |
| *GCA_009945535* | Mycoplasma pneumoniae | 2104 |
| *GCA_009945865* | Mycoplasma pneumoniae | 2104 |
| *GCA_009946285* | Mycoplasma pneumoniae | 2104 |
| *GCA_009946845* | Mycoplasma pneumoniae | 2104 |
| *GCA_009947205* | Mycoplasma pneumoniae | 2104 |
| *GCA_009947575* | Mycoplasma pneumoniae | 2104 |
| *GCA_009947985* | Mycoplasma pneumoniae | 2104 |
| *GCA_009948395* | Mycoplasma pneumoniae | 2104 |
| *GCA_009975385* | endosymbiont DhMRE of Dentiscutata heterogama | 1609546 |
| *GCA_012516495* | Mycoplasma sp. 1654_15 | 2725994 |
| *GCA_012934855* | Mycoplasma sp. Phocoena C-264-GEN | 754517 |
| *GCA_012934885* | Mycoplasma sp. C264-NAS | 2726117 |
| *GCA_013348745* | Mycoplasma sp. OR1901 | 2742195 |
| *GCA_014352955* | Mycoplasma sp. Pen4 | 640330 |
| *GCA_014803855* | Mycoplasma sp. | 2108 |
| *GCA_016925555* | Mycoplasma sp. Zaradi2 | 547987 |
| *GCA_017389835* | Mycoplasma sp. | 2108 |
| *GCA_900476125* | Mycoplasma alkalescens | 45363 |
| *GCA_900476175* | Mycoplasma putrefaciens | 2123 |
| *GCA_900660445* | Mycoplasma salivarium | 2124 |
| *GCA_900660465* | Mycoplasma pneumoniae | 2104 |
| *GCA_900660505* | Mycoplasma dispar | 86660 |
| *GCA_900660555* | Mycoplasma conjunctivae | 45361 |
| *GCA_900660715* | Mycoplasma arthritidis | 2111 |
| *GCA_900660735* | Mycoplasma cloacale | 92401 |
| *GCA_902712995* | Candidatus Mycoplasma haemohominis | 1494318 |
| *GCA_000006625* | Ureaplasma parvum serovar 3 (strain ATCC 700970) | 273119 |
| *GCA_000008205* | Mycoplasma hyopneumoniae (strain J / ATCC 25934 / NCTC 10110) | 262719 |
| *GCA_000008225* | Mycoplasma hyopneumoniae (strain 7448) | 262722 |
| *GCA_000008245* | Mycoplasma synoviae (strain 53) | 262723 |
| *GCA_000008365* | Mycoplasma mobile (strain ATCC 43663 / 163K / NCTC 11711) | 267748 |
| *GCA_000008405* | Mycoplasma hyopneumoniae (strain 232) | 295358 |
| *GCA_000011225* | Mycoplasma penetrans (strain HF-2) | 272633 |
| *GCA_000011445* | Mycoplasma mycoides subsp. mycoides SC (strain PG1) | 272632 |
| *GCA_000012765* | Mycoplasma capricolum subsp. capricolum (strain California kid / ATCC 27343 / NCTC 10154) | 340047 |
| *GCA_000019345* | Ureaplasma parvum serovar 3 (strain ATCC 27815 / 27 / NCTC 11736) | 505682 |
| *GCA_000020065* | Mycoplasma arthritidis (strain 158L3-1) | 243272 |
| *GCA_000021265* | Ureaplasma urealyticum serovar 10 (strain ATCC 33699 / Western) | 565575 |
| *GCA_000025365* | Mycoplasma gallisepticum (strain R(high / passage 156)) | 710128 |
| *GCA_000025385* | Mycoplasma gallisepticum (strain F) | 708616 |
| *GCA_000025845* | Mycoplasma crocodyli (strain ATCC 51981 / MP145) | 512564 |
| *GCA_000085865* | Mycoplasma hominis (strain ATCC 23114 / NBRC 14850 / NCTC 10111 / PG21) | 347256 |
| *GCA_000169535* | Ureaplasma urealyticum serovar 8 str. ATCC 27618 | 626095 |

| | | |
|---|---|---|
| *GCA_000169555* | Ureaplasma urealyticum serovar 4 str. ATCC 27816 | 550748 |
| *GCA_000169575* | Ureaplasma urealyticum serovar 7 str. ATCC 27819 | 519849 |
| *GCA_000169595* | Ureaplasma urealyticum serovar 9 str. ATCC 33175 | 550773 |
| *GCA_000169895* | Ureaplasma parvum serovar 6 str. ATCC 27818 | 519851 |
| *GCA_000169915* | Ureaplasma urealyticum serovar 5 str. ATCC 27817 | 518603 |
| *GCA_000169935* | Ureaplasma urealyticum serovar 11 str. ATCC 33695 | 526347 |
| *GCA_000169955* | Ureaplasma urealyticum serovar 12 str. ATCC 33696 | 550747 |
| *GCA_000171355* | Ureaplasma parvum serovar 14 str. ATCC 33697 | 515609 |
| *GCA_000171375* | Ureaplasma parvum serovar 1 str. ATCC 27813 | 515608 |
| *GCA_000171395* | Ureaplasma urealyticum serovar 13 str. ATCC 33698 | 515611 |
| *GCA_000171555* | Ureaplasma urealyticum serovar 2 str. ATCC 27814 | 626096 |
| *GCA_000179035* | Mycoplasma suis (strain Illinois) | 768700 |
| *GCA_000203215* | Mycoplasma suis (strain KI_3806) | 708248 |
| *GCA_000209735* | Mycoplasma fermentans (strain ATCC 19989 / NBRC 14854 / NCTC 10117 / PG18) | 496833 |
| *GCA_000211295* | Mycoplasma hyorhinis (strain MCLD) | 936139 |
| *GCA_000211545* | Mycoplasma gallisepticum S6 | 1006581 |
| *GCA_000218525* | Mycoplasma ovipneumoniae SC01 | 997765 |
| *GCA_000224105* | Mycoplasma putrefaciens (strain ATCC 15718 / NCTC 10155 / C30 KS-1 / KS-1) | 743965 |
| *GCA_000227355* | Mycoplasma iowae 695 | 1048830 |
| *GCA_000238995* | Mycoplasma haemocanis (strain Illinois) | 1111676 |
| *GCA_000241125* | Mycoplasma hyorhinis GDL-1 | 1129369 |
| *GCA_000253075* | Mycoplasma mycoides subsp. capri LC str. 95010 | 862259 |
| *GCA_000253095* | Mycoplasma leachii (strain 99/014/6) | 866629 |
| *GCA_000277795* | Mycoplasma wenyonii (strain Massachusetts) | 1197325 |
| *GCA_000281235* | Mycoplasma haemolamae (strain Purdue) | 1212765 |
| *GCA_000283755* | Mycoplasma pneumoniae 309 | 1112856 |
| *GCA_000286675* | Mycoplasma gallisepticum VA94_7994-1-7P | 1159197 |
| *GCA_000286695* | Mycoplasma gallisepticum NC95_13295-2-2P | 1159198 |
| *GCA_000319365* | Candidatus Mycoplasma haemominutum 'Birmingham 1' | 1116213 |
| *GCA_000367185* | Mycoplasma flocculare ATCC 27716 | 1004152 |
| *GCA_000376625* | Mycoplasma putrefaciens Mput9231 | 1292033 |
| *GCA_000380285* | Mycoplasma yeatsii 13926 | 1188240 |
| *GCA_000383515* | Mycoplasma hyorhinis ATCC 17981 | 634996 |
| *GCA_000385075* | Mycoplasma hominis ATCC 27545 | 1267000 |
| *GCA_000385095* | Mycoplasma synoviae ATCC 25204 | 1267001 |
| *GCA_000387745* | Mycoplasma pneumoniae 19294 | 1213463 |
| *GCA_000400855* | Mycoplasma hyopneumoniae 168-L | 1116211 |
| *GCA_000420105* | Mycoplasma orale ATCC 23714 | 1266996 |
| *GCA_000420225* | Mycoplasma moatsii ATCC 27625 | 1278300 |
| *GCA_000427215* | Mycoplasma hyopneumoniae 7422 | 754503 |

### Supplementary material 1B Random Forest classification of host and tissue isolation-site

*Table 1 Classifier scoring using all or only 12 features for host and tissue classification*

| | SCORE HOST CLASSIFIER ALL FEATURES | SCORE HOST CLASSIFIER 12 FEATURES | SCORE TISSUE CLASSIFIER ALL FEATURES | SCORE TISSUE CLASSIFIER 12 FEATURES |
|---|---|---|---|---|
| **PRECISION** | 0.97 | 0.94 | 0.84 | 0.89 |
| **RECALL** | 0.97 | 0.89 | 0.84 | 0.88 |
| **FSCORE** | 0.97 | 0.91 | 0.84 | 0.88 |

#### Hyper parameter optimization

**Hyper parameter search grid (searched with oob_score = True and 10x cross validation):**

{'n_estimators': [200, 400, 600, 800, 1000, 1200, 1400, 1600, 1800, 2000], 'max_features': ['auto', 'sqrt'], 'max_depth': [10, 20, 30, 40, 50, 60, 70, 80, 90, 100, 110, None], 'min_samples_split': [2, 5, 10], 'min_samples_leaf': [1, 2, 4], 'bootstrap': [True, False]}

**Optimal hyper parameters host classifier using all features:**

{'n_estimators': 1600, 'min_samples_split': 5, 'min_samples_leaf': 1, 'max_features': 'auto', 'max_depth': 10, 'bootstrap': True}

**Optimal hyper parameters host classifier using12 most important features:**

{'n_estimators': 1200, 'min_samples_split': 6, 'min_samples_leaf': 3, 'max_features': 'sqrt', 'max_depth': None, 'bootstrap': True}

**Optimal hyper parameters tissue classifier using all features:**

{'n_estimators': 400, 'min_samples_split': 5, 'min_samples_leaf': 1, 'max_features': 'sqrt', 'max_depth': 30, 'bootstrap': True}

**Optimal hyper parameters tissue classifier using 12 most important features:**

{'n_estimators': 400, 'min_samples_split': 6, 'min_samples_leaf': 2, 'max_features': 'sqrt', 'max_depth': 1, 'bootstrap': True}

#### Confusion matrixes

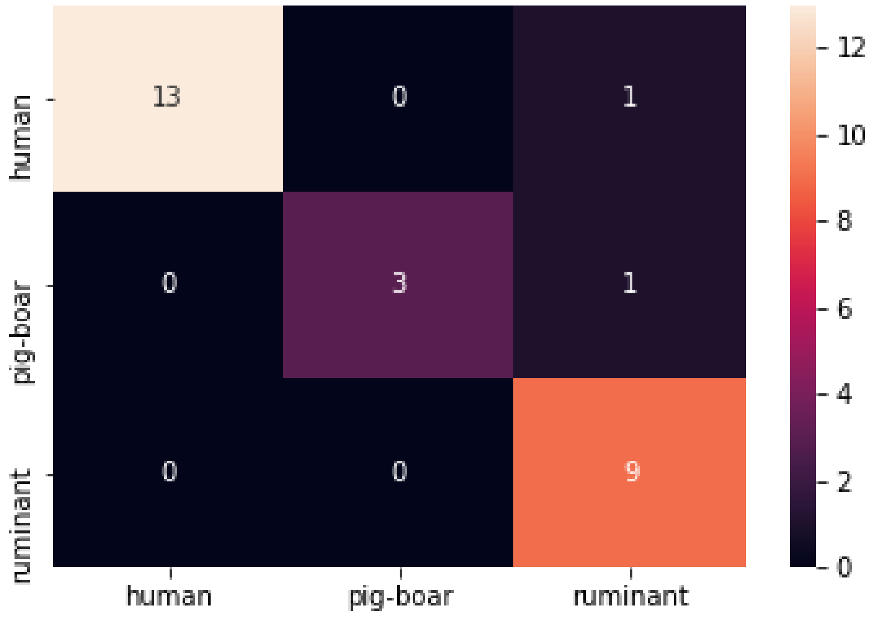

*Figure 1 Confusion matrix of the host classifier using the top 12 most important features*

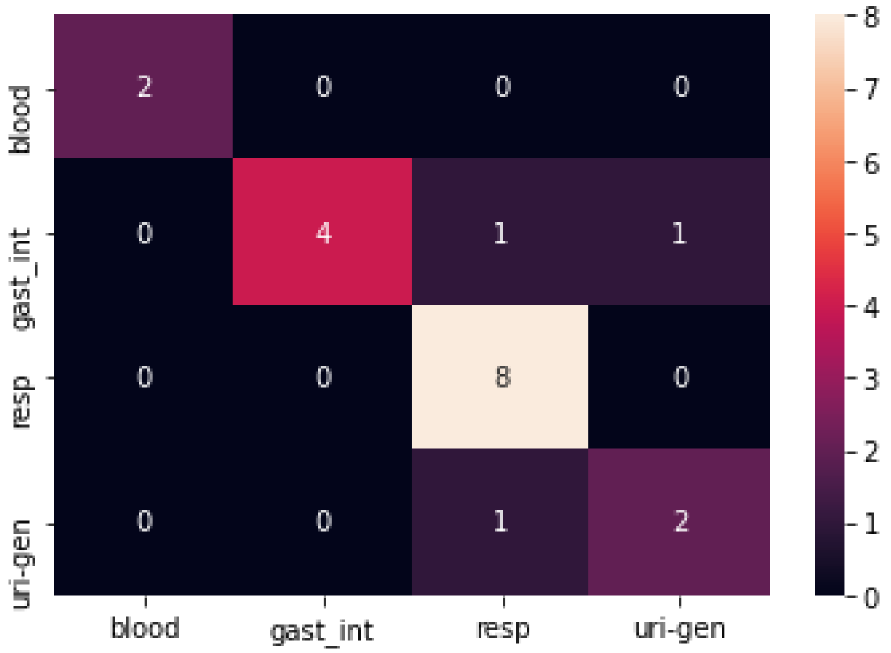

*Figure 2 Confusion matrix of the tissue classifier using the top 12 most important features*

#### Results for dual host_tissue classification

Only 52 genomes have both host and tissue information, predominantly human infecting Mycoplasma. Trained model completely failed to predict the test data, which only contained 2 out of 4 classes, '*human_gast_int*', '*human_resp'.* In conclusion, insufficient data as well as possibly to much similarity between the two classes *'human_gast_in*t', '*human_resp'*. Still the results are surprising since none of the genomes were correctly classified.

Test Accuracy base model on train and test data:: 1.0,0.0
Test Accuracy BEST model on train and test data:: 0.9743589743589743,0.0
Test f1_score BEST model on train and test data:: 0.9686520376175549,0.0
